# Evidence of compensation in the brain networks of Lewy body dementia and Alzheimer’s disease patients

**DOI:** 10.1101/159491

**Authors:** Luis R. Peraza, Ruth Cromarty, Xenia Kobeleva, Michael J. Firbank, Alison Killen, Sara Graziadio, Alan J. Thomas, John T. O’Brien, John-Paul Taylor

## Abstract

Dementia with Lewy bodies (DLB) and Alzheimer’s disease (AD) require differential management despite presenting with symptomatic overlap. A human electrophysiological difference is a decrease of dominant frequency (DF) −the highest power frequency between 4-15Hz– in DLB; a characteristic of Parkinsonian diseases. We analysed electroencephalographic (EEG) recordings from old adults: healthy controls (HCs), AD, DLB and Parkinson’s disease dementia (PDD) patients. Brain networks were assessed with the minimum spanning tree (MST) within six EEG bands: delta, theta, high-theta, alpha, beta and DF. Patients showed lower alpha band connectivity and lower DF than HCs. Lewy body dementias showed a randomised MST compared with HCs and AD in high-theta and alpha but not within the DF. The MST randomisation in DLB and PDD reflects decreased brain efficiency as well as impaired neural synchronisation. However, the lack of network topology differences at the DF indicates a compensatory response of the brain to the neuropathology.

## Introduction

Dementia with Lewy bodies (DLB) and Alzheimer’s disease (AD) are leading causes of neurodegenerative dementia in the elderly population. DLB is characterised by the core symptoms of visual hallucinations, cognitive fluctuation and Parkinsonism. Other symptoms may also be present and precede the core ones such as autonomic dysfunction, falls, and sleep disturbances (1). In contrast, patients with AD compared to DLB often have more significant memory problems; including memory loss, difficulty recalling recent events and learning new information (2). At early stages both dementias present with an important symptomatic overlap (3), e.g. problems with thinking and reasoning, leading to a frequent misdiagnosis of DLB patients as AD (4). Another common cause of dementia is Parkinson’s disease dementia (PDD); this condition shares symptomatic and pathologic overlaps with DLB (5). However their clinical paths differ at the beginning of the disease; in DLB motor symptoms either occur concurrently with cognitive symptom onset or after, whereas PDD patients experience motor symptoms before the onset of cognitive decline (1). AD and PDD offer two different disease perspectives for the understanding of DLB and this reflects the relative loading of pathology between the conditions, with DLB having both Alzheimer and alpha-synuclein pathology (6).

Electroencephalography (EEG) represents an important neuroimaging modality to study the effects of dementia in brain processes; previous work on this field has been carried out by Bonanni, Perfetti (7), Stam, de Haan (8) and Babiloni, Del Percio (9) amongst others (10, 11), with the current consensus indicating that there is a slowing of alpha band frequency, mainly at occipital regions in Lewy body dementia and that this feature may be useful as a diagnostic biomarker of DLB (12). Compared with other neuroimaging modalities, EEG has the advantage of being non-invasive, widely available and inexpensive. Additionally, EEG can be acquired multiple times without associated side effects (13).

The dominant frequency (DF) is perhaps the most promising EEG feature used to differentiate DLB from AD and it is defined as the frequency with the highest EEG power at the vicinity of the alpha band (7). In this regard Bonanni, Thomas (14) utilised compressed spectral arrays and showed a decrease of DF in DLB patients in occipital cortices. Other proposed features which are gaining traction include that of brain networks (15), which offer an integrative perspective for the complex functional connectivity of the brain and of particular interest is the subnetwork known as the minimum spanning tree (MST) (16). A brain network is comprised of nodes and edges; in EEG the nodes are the electrodes and the edges are the connectivity between electrodes. Network connectivity can be estimated with the phase lag index (PLI), a connectivity measure resistant to the effects of the scalp’s volume conduction (17). The MST is then the network which has the minimum number of strongest edges while connecting all nodes without cycling paths. Using this approach, Yu, Gouw (18) reported that in AD there exists a decentralisation of the network tree towards a line-like less efficient configuration. In the DLB field, van Dellen, de Waal (19) showed that patients present with a decreased EEG network efficiency, which the authors linked to impaired cognition.

In this investigation we studied brain network connectivity in EEG recordings from DLB, AD, PDD, and healthy control (HC) participants in the sensor domain using the MST. We hypothesised that network structural features and their variability consistently differ among our studied dementia groups and that these may have potential use as differential biomarkers in DLB diagnosis. Specifically, we hypothesised that the functional network topology of our patient groups in the DF band would highlight pathological mechanisms of dementia and neural degeneration.

## Results

After EEG pre-processing, 6 AD, 1 HC, 1 DLB and 1 PDD participants were excluded from the final analysis because their cleaned EEG recordings had less than 50 seconds of continuous EEG, resulting in final participant groups of 17 HC, 26 AD, 25 DLB and 21 PDD participants.

### Demographics

Demographics and clinical variables of our studied participant groups are given in Table 1. The recruited groups did not show significant differences in terms of age and gender. All patient groups were matched for global cognitive impairment by their CAMCOG and MMSE scores, although the AD showed a trend of lower cognitive impairment compared with the other dementia groups. The memory domain in the CAMCOG was significantly lower in the AD group than in PDD and DLB. Complex visual hallucinations (NPI hall) and cognitive fluctuations (CAF) were, as expected, higher in both Lewy body dementias compared to AD patients. In relation to their medications, PDD patients were on higher doses of dopaminergic therapy than the DLB group, but there were not significant differences between the dementia groups in terms of use of cholinesterase inhibitor medications.

**Table 1.** Demographics and clinical variables

|  | HC (N=17) |  | AD (N=26) |  | DLB (25) |  | PDD (21) |  | p-value |
| --- | --- | --- | --- | --- | --- | --- | --- | --- | --- |
| Age | 76.18 | ±5.65 | 76.46 | ±7.83 | 75.8 | ±6.5 | 73.29 | ±5.842 | F(3,85)=1.03, p-value=0.380<sub>⊥</sub> |
| Male/Female | 10/7 | | 18/8 | | 20/5 | | 19/2 | | $\chi^2(3, N=89)=5.89$, p-value=0.117<sub>⊥</sub> |
| MMSE | 29.12 | ±0.857 | 20.08 | ±4.279 | 22.64 | ±4.3 | 22.71 | ±4.85 | F(2,69)=2.81; p-value=0.067* |
| CAMCOG total | 96.47 | ±3.69 | 66.54 | ±16.46 | 76.08 | ±12.92 | 73.38 | ±13.88 | F(2,69)=2.89; p-value=0.063* |
| CAMCOG executive | 22.71 | ±2.25 | 14.46 | ±5.49 | 13.12 | ±5.03 | 12.9 | ±2.86 | F(2,69)=0.79; p-value=0.457* |
| CAMCOG memory | 23.47 | ±2.035 | 10.88 | ±5.1 | 18.04 | ±4.9 | 17.48 | ±5.05 | F(2,69)=15.5; p-value<0.001* |
| CAMCOG attention | 6.76 | ±0.562 | 3.81 | ±2.514 | 4.4 | ±2.27 | 4.19 | ±1.721 | F(2,69)=0.465; p-value=0.63* |
| NPI hall | 0 | 0 | 0.04 | ±0.2 | 1.76 | ±1.86 | 2.57 | ±2.29 | F(2,68)=14.1; p-value<0.001* |
| CAF total | 0 | 0 | 0.6 | ±1.52 | 3.92 | ±4.142 | 6.63 | ±4.271 | t(42)=2.12, p-value=0.03<sub>⊥</sub> |
| Animal<br>Categorical<br>fluency | 20.71 | ±5.71 | 11.15 | ±5.25 | 11.16 | ±3.88 | 11.33 | ±4.14 | F(2,69)=0.011; p-value=0.989* |
| UPDRS III | 1.24 | ±1.522 | 2.28 | ±1.929 | 15.52 | ±8.2 | 25.57 | ±7.018 | t(44)=4.42, p-value<0.001<sub>⊥</sub> |
| Angle<br>discrimination | 19.63 | ±0.88 | 18.24 | ±3.72 | 16.17 | ±4.85 | 16.55 | ±4.99 | H(2)=4.1, p-value=0.129<sub>⊥</sub> |
| FAS<br>Verbal fluency | 45.12 | ±16.53 | 25.96 | ±16.14 | 19.68 | ±12 | 20.81 | ±13.7 | H(2)=1.94, p-value=0.378<sub>⊥</sub> |
| LEDD | 0 | 0 | 0 | 0 | 164.88 | 231.56 | 818.23 | 381.26 | t(44)=7.15, p-value < 0.001<sub>⊥</sub> |
| ACHel<br>(yes/no) | 0/17 | | 24/2 | | 23/2 | | 16/4† | | $\chi^2(2, N=71)=2.13$, p-value=0.346 |

\* ANOVA three groups (AD, DLB, PDD);
⊥ ANOVA four groups;
⊥ $\chi^2$ test four groups
⊥ Unpaired t-test (DLB vs PDD);
⊥ Kruskal-Wallis three groups (AD, DLB, PDD).
† One PDD patient was on Memantine

### Dominant frequency and network measures

The one-way ANOVA tests were significant for differences in DF across the four participant groups; F(85,3)=6.75, p-value=0.0004. Post-hoc tests showed that DLBs had a significantly lower DF (DLB: mean = 6.8Hz, SD = 0.91) than ADs (mean = 7.5Hz, SD =1.21) and HCs (mean=8.6Hz, SD=0.82) but not when compared against the PDD group (PDD: mean=6.2Hz, SD=0.56). The AD group showed a lower DF than the HCs but it was higher than in the PDD group, Figure 1. The one-way ANOVA for the dominant frequency variability (DFV) also showed significant differences; F(85,3)=4.59, p-value=0.005. The AD group presented with significantly higher variability (AD: mean=1.67Hz, SD=0.79) compared with the other groups, and DLBs showed significantly lower DFV (DLB: mean=1.027Hz, SD=0.3) when compared with AD, Figure 1.

**Figure 1.**
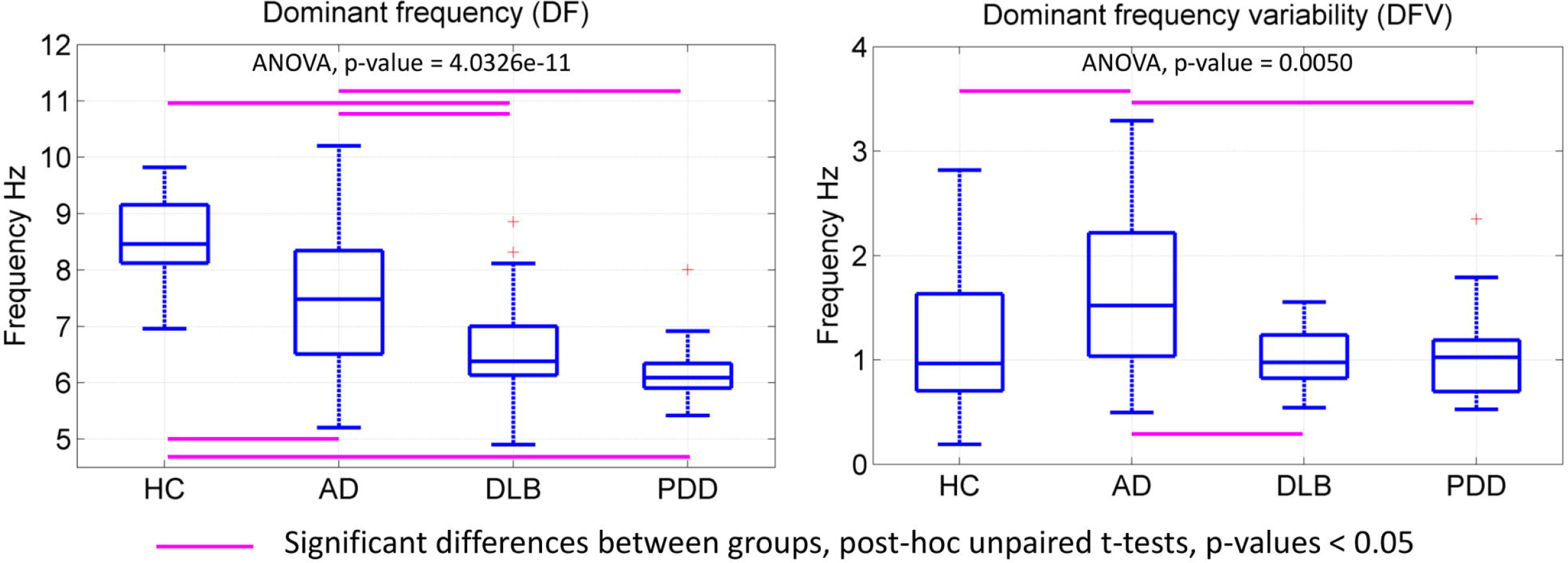
Dominant frequency (DF) and variability (DFV) box-plots. Four-group ANOVAs’ p-values are given for each test, uncorrected. DF mean and standard deviation (SD) per group: HC (8.59Flz, SD=0.82), AD (7.5Hz, SD=1.21), DLB (6.8Hz, SD=0.91), PDD (6.2Hz, SD=0.56). DFV mean and SD per group: HC (1.21Hz, SD=0.67), AD (1.67Hz, SD=0.79), DLB (1.027Hz, SD=0.3), PDD (1.048Hz, SD=0.42). Each box in the plots shows the median as the central line, the extremes of each box are the first and third quartile and the whiskers represent the minimum and maximum values in the sample. DF and DFV were estimated from the occipital electrodes; see Figure 1-Figure Supplement 1.

The two-way four-group ANOVA tests (group and frequency band effects) for the network structure measures within the six frequency bands, showed significant differences for the maximum nodal degree (degree_max_), leaf ratio, eccentricity, network diameter and radius; these measures were significant for the group effect. The global effects for these measures showed a decreased degree_max_ and leaf ratio in the three dementia groups compared with HCs, predominantly in the alpha band, while diameter, eccentricity and radius were higher in the dementia groups compared with HCs, predominantly in the high-theta band and with a trend for differences in the means of HC < AD < DLB < PDD, see Figure 2 upper row. However, when analysing the multiple post-hoc between-group differences in network topology measures within the DF band, none of the post-hoc tests resulted significant showing uncorrected p-values > 0.05, Figure 2 bottom row.

**Figure 2.**
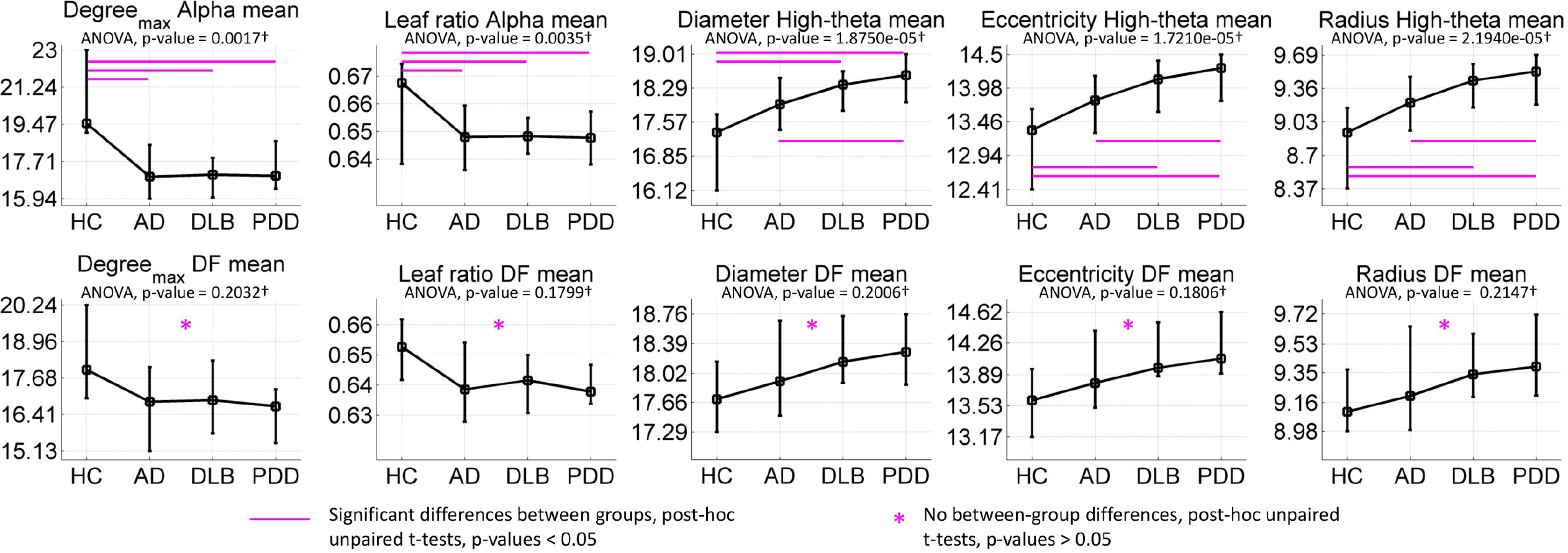
Two-way four-group ANOVA tests for the group effect. Upper row: The most significant post-hoc one-way ANOVAs per network measure and frequency band are shown. Significant (at p-value < 0.05 for the multiple unpaired t-tests and uncorrected) post-hoc between-group differences for the group means are indicated with line markers in magenta colour. Bottom row: post-hoc oneway ANOVA tests for the dominant frequency (DF) per network measure were nonsignificant. *Post-hoc multiple unpaired t-tests for comparisons between participant groups were also nonsignificant for DF. +P-values are shown uncorrected for multiple comparisons. For each plot, the line crosses the group means and the error bars span the first and third quartiles. Group effects are given in Figure 2-Table Supplement 1 and their mean in Figure 2-Table Supplement 2.

The MST PLI related measures (PLI mean, leaf, root and height) were significant for both mean and variability but mainly for the interaction effect of group and frequency band —see Figure 3. Frequencies at which differences between groups were found predominantly in the theta, high-theta and alpha bands. Specifically, the HC group was significantly different when compared against the dementia group for PLI measures in the alpha band. PLI height was significantly different between DLB and AD in the beta band.

**Figure 3.**
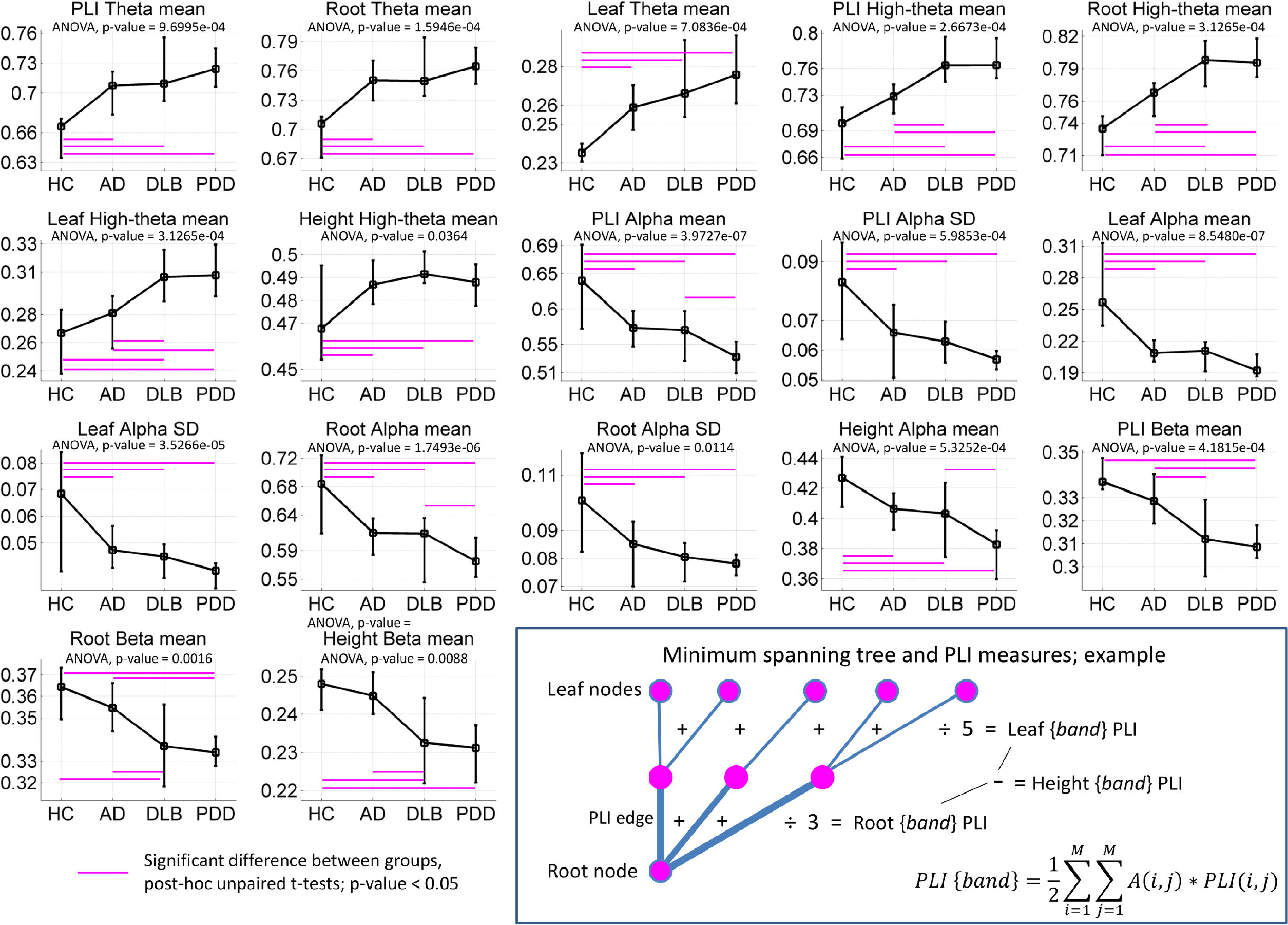
Minimum spanning tree measures that showed significant differences across the groups of interests; ANOVA tests’ p-values are shown Bonferroni adjusted, i.e. multiplied by a correction factor that accounts for the 72 post-hoc one-way ANOVA tests. Significant multiple unpaired t-tests, p-value < 0.05, within each one-way ANOVA test are shown with line markers (in magenta colour) for the two groups with different means. For each plot, the line crosses the group means and the error bars span the first and third quartiles; mean and standard deviation are given in Figure 3-Table Supplement 2. Bottom-right: Minimum spanning tree example comprising nine nodes and eight edges, where five of them connect leaf nodes. *A(i,j)* is the network adjacency matrix where an edge between nodes *i* and *j* is indicated with 1, and 0 otherwise. *PLI(i,j)* is the PLI score between nodes / and *j. M* is the total number of electrode nodes. See Figure 3-Table Supplement 1 for variable meaning.

Significant multiple linear regressions between the clinical variables and the network measures are shown in Table 2 and Figure 4. None of the measures of variability related with the clinical variables, and we also tested these without the log-transformation with same results. For dominant frequency, the most significant relation was found between it and verbal fluency in all dementia groups with a positive correlation, and in AD and DLB for the within-group correlations, see Table 2. Dominant frequency was also significantly related to animal naming, CAMCOG, MMSE and Trail A test, and showed a positive relation between DF and cognitive impairment by these clinical measures. PLI tree measures in the theta band significantly related to cognitive variables. The network mean PLI in the high-theta band significantly related to the frequency and severity of complex visual hallucinations (NPI hall) in the DLB group, see Figure 4. We also tested whether changing the reference group in the multiple linear regressions altered the results, which it did not.

**Figure 4.**
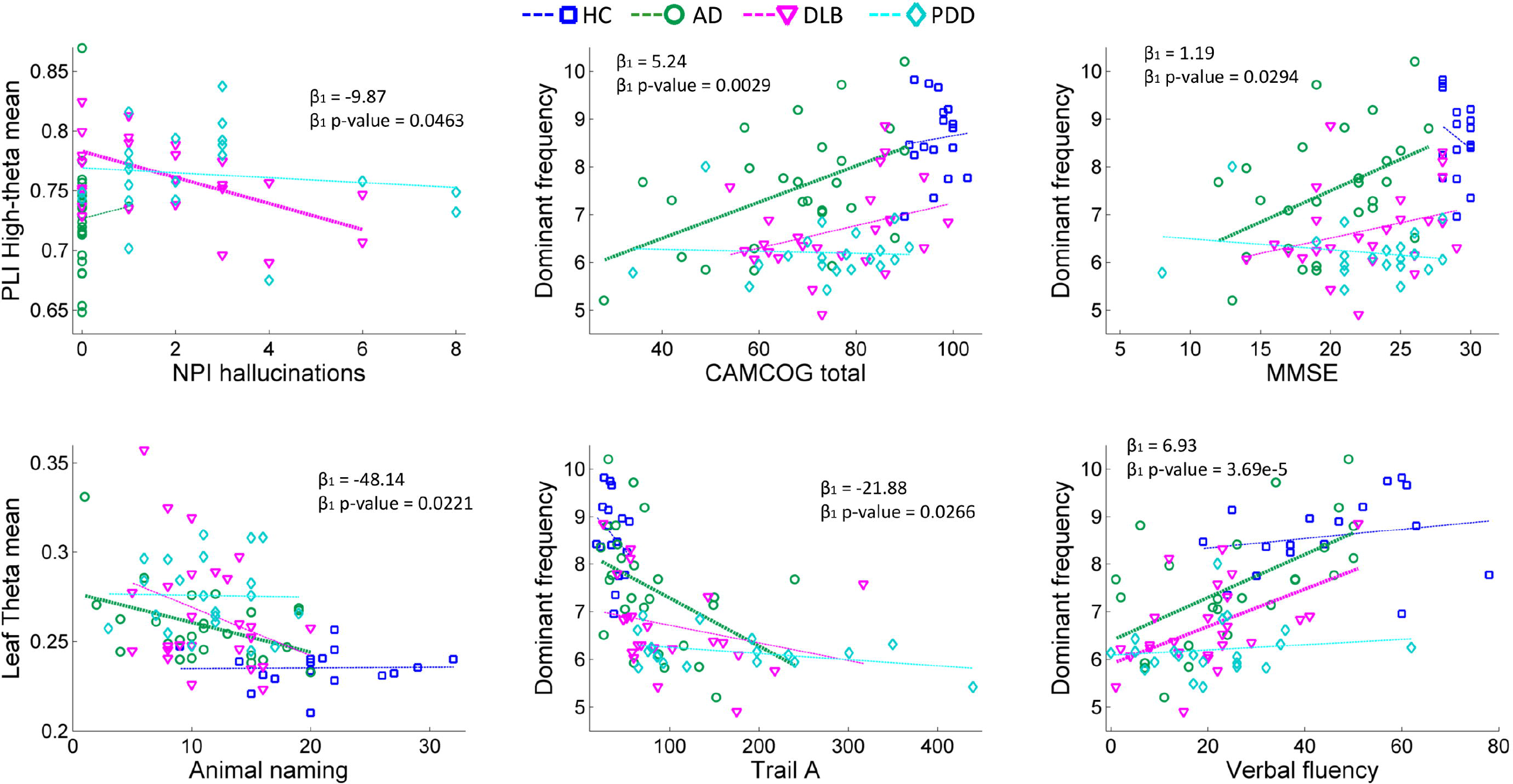
Selected significant multiple liner regressions between clinical variables and network measures. *P-values for the β_1_ coefficient for network measure are shown uncorrected. Significant within group Pearson’s correlation, at p-value < 0.05, are shown with thicker dashed lines, see Table 2 for all significant results. Model: Clinical variable ∼ β_1_*network variable + β_2_*D_DLB_ + β_2_*D_pdd_, where D is a dichotomous vector for each of the Lewy body dementia groups.

**Table 2.** Multiple linear regressions controlling for group membership with two dichotomous variables (D) for the Lewy body dementia groups. Model: Clinical variable ∼ β_1_*network variable + β_2_*D_DLB_ + β_2_*D_PDD_- Shown p-values correspond to the β coefficient of the network variable in the regression model and are shown uncorrected. Within group Pearson’s correlation and p-values are also given; ρ (p-value) and full model’s coefficients are given in Table 2-1.

| Network variable | Clinical variable | Linear regression [AD+DLB+PDD] | | HC<br>$\rho$ (p-value) | AD<br>$\rho$ (p-value) | DLB<br>$\rho$ (p-value) | PDD<br>$\rho$ (p-value) |
| --- | --- | --- | --- | --- | --- | --- | --- |
| | | $B_1$ | $B_1$ p-value | | | | |
| PL high-theta mean | NPI hallucinations | -9.87 | 0.0463 | – (–) | 0.047 (0.824) | **-0.498 (0.011)** | -0.127 (0.584) |
| PLI theta mean | Animal naming | -31.47 | 0.0400 | 0.22 (0.398) | -0.36 (0.075) | -0.31 (0.129) | 0.05 (0.842) |
| PLI theta mean | CAMCOG memory | -39.83 | 0.0204 | -0.04 (0.866) | -0.23 (0.261) | -0.31 (0.137) | -0.33 (0.147) |
| PLI theta mean | CAMCOG total | -119.70 | 0.0153 | 0.06 (0.834) | -0.33 (0.105) | -0.32 (0.116) | -0.21 (0.361) |
| Leaf theta mean | Animal naming | -48.14 | 0.0221 | 0.02 (0.938) | **-0.43 (0.027)** | -0.32 (0.120) | -0.02 (0.933) |
| Leaf theta mean | CAMCOG memory | -58.91 | 0.0125 | 0.19 (0.460) | -0.31 (0.130) | -0.28 (0.174) | -0.36 (0.113) |
| Leaf theta mean | CAMCOG total | -169.77 | 0.0124 | 0.21 (0.430) | -0.37 (0.061) | -0.33 (0.110) | -0.21 (0.373) |
| Root theta mean | CAMCOG memory | -43.24 | 0.0196 | -0.03 (0.914) | -0.17 (0.395) | -0.29 (0.148) | -0.43 (0.054) |
| Root theta mean | CAMCOG total | -126.38 | 0.0178 | 0.07 (0.788) | -0.25 (0.220) | -0.32 (0.124) | -0.32 (0.161) |
| Dominant frequency | Animal naming | 1.36 | 0.0131 | 0.3 (0.244) | **0.41 (0.040)** | 0.25 (0.230) | 0.03 (0.902) |
| Dominant frequency | CAMCOG attention | 0.59 | 0.0317 | -0.27 (0.292) | 0.34 (0.088) | 0.38 (0.062) | -0.38 (0.091) |
| Dominant frequency | CAMCOG executive | 1.54 | 0.0071 | 0.22 (0.393) | 0.38 (0.056) | 0.32 (0.124) | 0.02 (0.918) |
| Dominant frequency | CAMCOG total | 5.24 | 0.0029 | 0.08 (0.756) | **0.5 (0.010)** | 0.33 (0.104) | -0.05 (0.819) |
| Dominant frequency | MMSE | 1.19 | 0.0294 | -0.25 (0.336) | **0.45 (0.021)** | 0.30 (0.141) | -0.2 (0.389) |
| Dominant frequency | Trail A | -21.88 | 0.0266 | -0.28 (0.270) | **-0.42 (0.034)** | -0.27 (0.220) | -0.39 (0.112) |
| Dominant frequency | Verbal fluency | 6.93 | 3.69E-05 | 0.20 (0.439) | **0.59 (0.002)** | **0.52 (0.008)** | 0.16 (0.559) |

### Diagnostic analysis

MST measures that showed significant differences for the diagnostic scenarios including DF and DFV, were included in the classification analysis; the receiver operating characteristic curves for this analysis are shown in Figure 5. AD vs DLB classification reached an 80% sensitivity (0.41-0.82 95% confidence internal CI), 85% specificity (0.42-0.92 CI) with 0.86 area under the curve (AUC). When differentiating DLB versus PDD, sensitivity was 76% (0.27-0.83 CI) and specificity was also 76% (0.58–0.88 CI); 0.77 AUC. Classification between HC and AD+DLB reached 0.96% sensitivity (0.78-1.0 CI) and 77% specificity (0.58-0.90 CI), 0.93 AUC. The step-wise multiple linear regression showed that for the AD vs DLB separation, the DFV was the best discriminant between both dementias (p-value = 0.0083) followed by PLI height in the beta band (p-value = 0.027). When diagnosing both combined groups AD+DLB versus the HC group, the step wise linear regression showed that the best discriminants were the PLI leaf mean in the alpha band (p-value < 0.0001) followed by DF (p-value < 0.0039). For the classification between DLB and PDD the PLI mean in the alpha band was the best discriminant between both groups (p-value = 0.0012).

**Figure 5.**
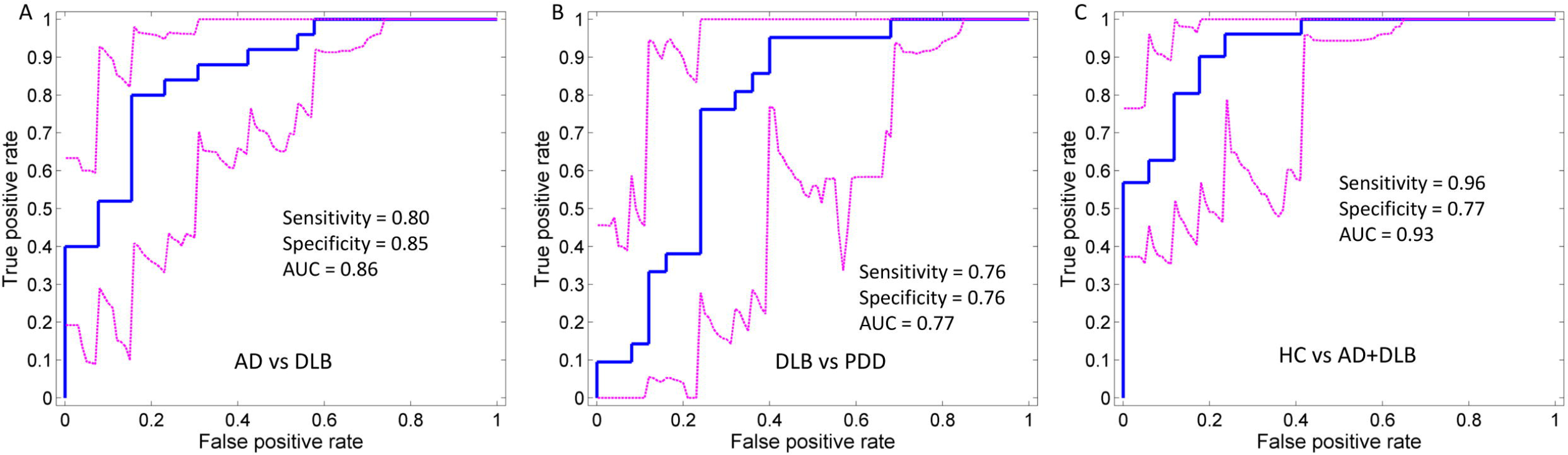
Receiver operating characteristic (ROC) curves by logistic regression for each of the diagnostic scenarios analysed, see Figure 5-Table Supplement 1. Magenta dashed lines show the 95% confidence intervals for the ROC curves; bootstrapping with 5000 iterations implemented with the perfcurve.m MATLAB function. Optimal sensitivity and specificity as well as the area under the curve (AUC) are shown within each curve plot.

## Discussion

Our analysis showed significant differences between the dementia groups and HCs for DF and PLI network measures. We confirmed the well-established finding that there is a slowing of DF in the Lewy body dementias −DLB and PDD– compared with AD and HCs in the occipital regions (9, 13, 20, 21). Additionally, we explored the variability of DF between the studied groups and we demonstrated that DFV is higher in AD compared with HCs and both Lewy body dementia groups. Differences between the MST measures were significant at several frequency bands, mainly for theta, high-theta and alpha. However, PLI height scores in the beta band was shown to be lower in DLB compared with AD patients. PLI height is a measure of focused synchronisation in the root node (high PLI and degree_max_) and to be higher requires the existence of leaves far away from the root with low PLI connections, these latter lower than the root node connections (22). A decrease in PLI height will therefore occur in two scenarios: 1) a star-like topology where the majority of edges of the root and leaf nodes are the same or 2) a PLI random connectivity matrix, where there are not significant differences in PLI strength between leaf edges and root edges. Our results suggests the latter case, i.e. that our DLB and PDD groups present with a randomisation of the MST. This is observed by the global effects of the network measures; the lower degree_max_ and leaf ratio in the alpha band combined with higher network diameter, eccentricity and radius in the high-theta band indicate network randomisation and a decrease in global efficiency of the MST. Similar results were reported by Utianski, Caviness (23) who showed than in PDD there exists a less “star-like” network topology or as Yu, Gouw (18) reported, a “line-like” topology. Visualisation of these MST network characteristics in four representative participants in our groups is shown in Figure 6, for the alpha and dominant frequency bands. The star-like topology can be easily observed in the HC participant where two highly connected nodes (with high nodal degree) comprise the centre of the star. The randomised tree or line-like topology can be observed easier in the DLB patient in the dominant frequency example (bottom row); nodes here are not highly connected (low degree_max_) and several nodes are connected within the same path in a sequence, i.e. lines.

**Figure 6.**
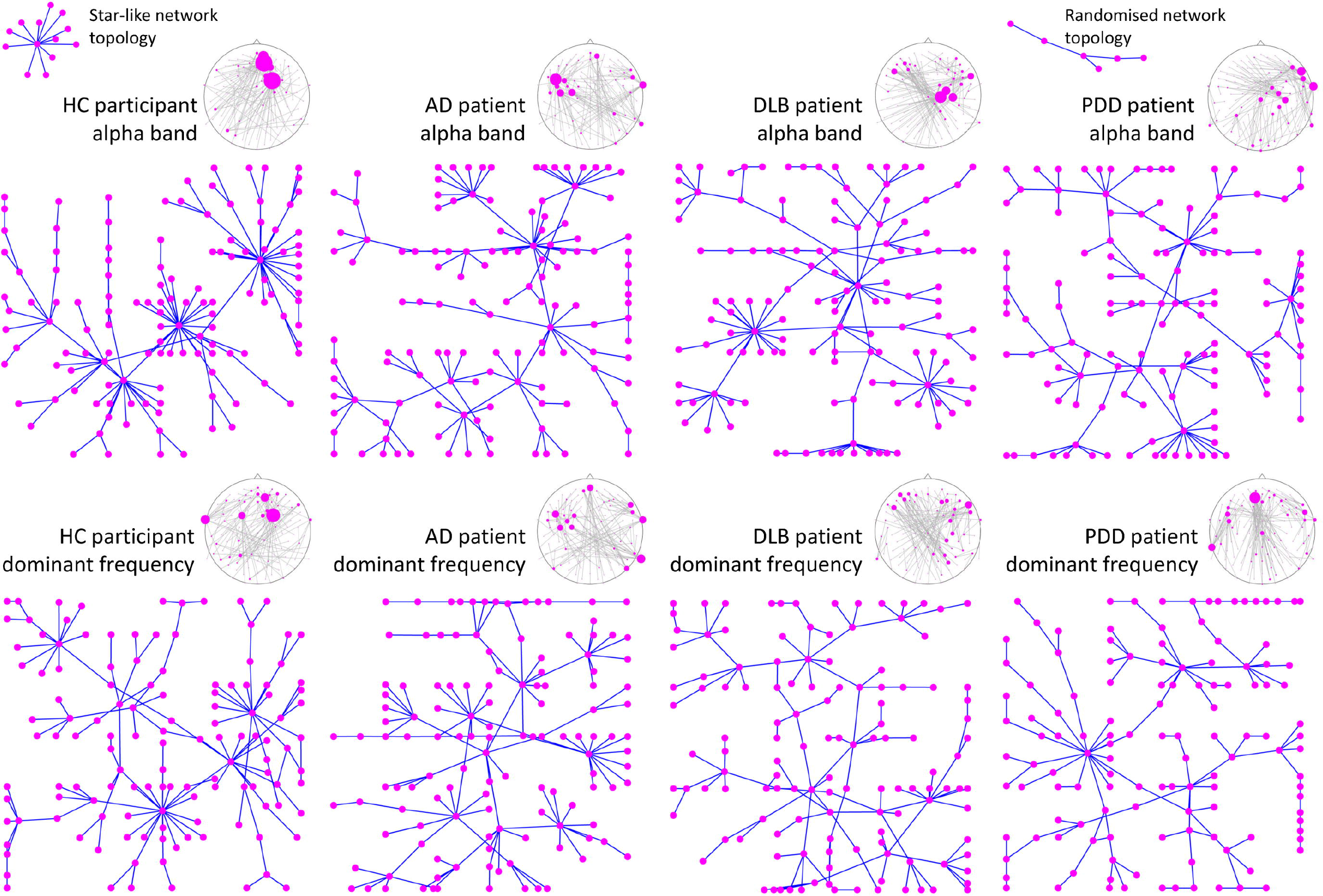
EEG minimum spanning tree examples from representative participants. Each participant was chosen as representative for being the median of their respective groups according to the leaf ratio mean in the alpha band. Similarly, the dominant frequency tree is also shown for the same participants. Notice the distinguishable “star-like” topology for the alpha-band network tree of the healthy control (HC) with two nodes with high nodal degree, and the more random tree in Lewy body dementia patients with a “line-like” topology; dementia with Lewy bodies (DLB) and Parkinson’s disease dementia (PDD). The Alzheimer’s disease (AD) patient also shows a star-like topology, although not as evident as the HC, suggesting impaired network. At the top of each network tree, the same network is shown on an EEG 10-5 system layout; here the diameter of the EEG electrodes is proportional to their nodal degree.

In a previous investigation, van Dellen, de Waal (19) reported a PLI connectivity decrease in the alpha band of DLB patients compared to AD and HCs. This contrasts somewhat with our investigation where PLI alpha was not significantly different between AD and DLB, although we demonstrated that it was lower in both dementias when compared with HCs. Instead we found that there are PLI differences in the high-theta and beta band between AD and DLB. PLI alpha band differences in our study occurred when comparing HCs against all dementia groups but not between the dementia groups. This suggests that PLI in the alpha band is a marker of healthy brain but that it is not specific enough to differentiate between dementias (9).

Exploratory diagnostic classification between AD and DLB was achieved by MST-PLI measures in the high-theta and beta bands as well as DF and DFV with high sensitivity and specificity of 80% and 85% respectively, and which is comparable to FP-CIT SPECT imaging (24). For this classification analysis, the most significant discriminant was the larger DFV in the AD group followed by the lower PLI height in beta band in the DLB group. This contrasts with the results by Bonanni, Thomas (14) who reported higher DFV in DLB patients compared to AD ones and both dementia groups showed higher DFV than HCs (14). This result disagreement may be driven by cohort specific differences or by the estimating method for DFV; here DFV was defined as the standard deviation of the DF throughout the EEG segments, while in Bonanni, Thomas (14) DFV was defined using a visual rating of DF range on sequential EEG segments. Forward projection of these MST-PLI and DF measures onto independent EEG samples will be required to clarify the robustness of these measures as diagnostic tools.

DF and MST-PLI frequency scores significantly related with cognitive variables in our patients, suggesting a pathological relation with dementia. The verbal fluency test, which measures executive function, was highly positively correlated with the dominant frequency in the dementia groups. Complex visual hallucinations (severity and frequency) was related with PLI mean in high-theta band, and significantly correlated within the DLB group, see Table 2. To date, we are not aware of previous investigations reporting a relation between complex visual hallucinations and resting state EEG features in DLB; this core symptom is frequently studied with event related responses (25, 26) or transcranial stimulations (27). However, previous EEG investigations in the Charles Bonnet syndrome (CBS) which is characterised by complex visual hallucinations in persons with partial or severe blindness, have reported alterations in alpha and theta power in CBS patients (28, 29) and this agrees with our results. Nevertheless, we must also take into account that our multiple linear regression results are uncorrected, and thus these findings must be taken with caution.

None of the MST measures were significantly different between groups for the individualised DF band in measures related to network structure such as diameter, radius and eccentricity; see Figure 2 bottom row. This suggests that in dementia, the EEG network which is present at lower dominant frequencies and whilst it is pathological, it is still a network which would have been present in the premorbid state with normal structural neural communications, and thus may represent a reactive or compensation state (30). Brain compensation is a phenomenon often reported in functional magnetic resonance imaging (fMRI) research (31). In fMRI, compensation is expressed and interpreted as an over-recruitment of neural circuits in order to maintain cognitive levels or motor processes in the presence of disease. In this regard, computational models of cortical activity have proven that the resonance frequency – dominant frequency– of neural circuits is inversely proportional to the size of the excited tissue (32), i.e. larger networks communicate at lower frequencies (33). These investigations thus suggest that the decrease of dominant frequency observed in our Lewy body dementia and Alzheimer’s disease participants may be an effect of overrecruitment of neural tissue in order to maintain dynamic network structure; or in other words, recruitment of larger neural circuits needed for compensation is implemented by the brain at lower frequencies. Confirmatory preliminary evidence from the same patients reported in the current study using functional task imaging, demonstrated increased fMRI-BOLD activations compared with healthy controls and which were related to task difficulty (34).

In conclusion, we investigated functional network differences between HCs and three dementia groups; AD, DLB and PDD. We found that the alpha band and its related MST network measures are markers of healthy brain as compared with dementia, while DLB and AD patients differentiated in DFV and PLI height in the beta band and these measures resulted of higher utility as potential diagnostic biomarkers. Also, we demonstrated that DLB as well as PDD, present with a randomised MST network compared to HCs and this suggests a decrease in focused neuronal synchronisation in these patients for the high-theta and alpha bands. In contrast, we found evidence of compensation in the DF functional network where no topological differences were found between our dementia groups and healthy controls for their MSTs.

## Methods

### Participants and clinical assessments

A total of 98 old adult participants were recruited within the North East of England. Dementia patients were recruited from a population referred to old age psychiatry and neurology services: 32 patients with AD (22 male, 10 female), 26 patients with DLB (21 male, 5 female), and 22 patients diagnosed with PDD (20 male, 2 female). Additionally, 18 healthy controls (HC) were recruited for group comparisons (11 male, 7 female). Patient groups were diagnosed by two experienced clinicians according to the current clinical diagnosis criteria for these dementias: The revised dementia with Lewy bodies consensus criteria (1), the diagnostic criteria for PDD (35), and the National Institute on Aging-Alzheimer’s Association criteria for AD (36).

All participants underwent comprehensive neurological and neuropsychiatric examinations. These were the Mini-mental state examination (MMSE), the Cambridge cognitive battery tests (CAMCOG), the Unified Parkinson’s Disease rating scale part III (UPDRS III), trail making test A, animal naming and visual perception (angle discrimination task). Additionally, the cognitive assessment of fluctuations (CAF)(37) for their frequency and duration, and the neuropsychiatric inventory test for the severity and frequency of hallucinations (NPI hallucinations) assessed; for the latter, carers were specifically asked about visual hallucinations rather than hallucinations in other modalities (38). For the patients that were on dopaminergic therapy, the Levodopa equivalent daily dose (LEDD) was estimated (39). A MMSE score < 12 for patients and < 26 for the HC group was used as exclusion criteria. None of the recruited healthy participants had a history of neurological or psychiatric conditions and neither did the patients outside their core dementia. Patients with Parkinsonism and on dopaminergic therapy were studied in the “ON” state. Ethical approval for this study was given by the Northumberland Tyne and Wear NHS Trust and Newcastle University ethics committee. All participants gave written informed consent prior study participation.

### Electroencephalography and signal processing

Resting EEG recordings were acquired with Waveguard caps (ANT Neuro, The Netherlands) comprising 128 sintered Ag/AgCl electrodes placed in a 10-5 positioning system (40). Channels were recorded with a sampling frequency of 1024 Hz and electrode impedance of < 5KΩ. At recording, all electrodes were referenced to Fz channel and the ground electrode was attached to the right clavicle. 150 seconds of continuous EEG resting state eyes closed were stored for off-line data processing, participants were seated throughout the recording and instructed to remain awake and as still as possible.

All pre-processing steps were carried out blinded to group membership and implemented with EEGLAB (41) MATLAB functions (R2012; MathWorks, Natick Massachusetts). 1) EEG channels had their baseline component subtracted and were band pass filtered between 0.3Hz and 54Hz with a second order Butterworth filter. 2) Bad channels were deleted including the reference electrode Fz; mean of four channels, with minimum one and maximums of 10 channels deleted. 3) Independent component analysis (ICA) with the FastICA (42) algorithm using default parameters was run for the entire EEG recording and up to 12 artefactual independent components were deleted. At this step special interest was put on identifying eye blink, cardiac and the 50Hz power line components, i.e. components that were noisy throughout the EEG recording. 4) If after component deletion artefactual components were still identified, FastICA algorithm was run again and up to 6 artefactual components were deleted. 5) Noisy EEG segments with artefacts affecting all channels (e.g. muscle artefacts) were deleted. 6) EEG components were remixed to the sensor domain. 7) Previously deleted bad channels were spatially interpolated with EEGLAB using spherical interpolation. 8) As final step, the electrode montage was modified to average montage (43). Participants with less than 50 seconds of continuous EEG after artefact cleaning were excluded from the remaining of the analysis.

### Dominant Frequency and its variability

Dominant frequency (DF) was estimated for the 50 second EEG segments. First, occipital channels (PO9, PO7, POO9h, PO5, O1, PO3, POO3h, OI1h, POz, Oz, PO4, POO4h, PO6, O2, OI2h, PO8, POO10h, PO10) were averaged to obtain a representative signal from this region (Figure 1-1). Power spectral density was then estimated with Welch’s periodogram; 2048 sample segments tapered with a Hamming window, 50% overlap between segments and a fast Fourier transform size of 2^^^13; these parameters led to a frequency resolution of 0.125Hz. We defined DF as the frequency bin in the power spectrum with highest power between 4 and 15Hz. DF for each participant was then estimated as the mean dominant frequency across the 49 windowed segments and DF variability (DFV) was defined as the standard deviation (SD) across the segments.

### Phase lag index

Phase lag index (PLI) was chosen to measure the regional relation among all electrode pairs (44). Briefly, PLI uses the Hilbert transform to estimate consistent causal delay between two signal sources and it has been proven that this measure is less affected by the scalp’s volume conduction, a problem with other measures such as spectral coherence or Pearson’s correlation (43). PLI scores are bounded between 0 and 1, where 0 means lack of causal synchronisation and 1 full causal synchronisation. In this investigation, PLI was estimated for six frequency bands: the standard EEG delta (0.5-4Hz), theta (4-5.5Hz), high-theta (5.5-8Hz), alpha (8-13Hz), beta (13-30Hz) and the dominant frequency band (DF ±2 Hz) whose centre was participant specific. To compute the PLI scores, the 50 second EEG recordings were first filtered at each of the defined frequency bands using a second order Butterworth band-pass filter, then Hilbert-transformed and finally segmented into two-second segments with one second overlap leading to a total of 49 segments. After this, 49 PLI connectivity matrices of dimension 128x128 elements were computed for each two-second segment, frequency band and participant.

### Minimum spanning tree

The minimum spanning tree (MST) in EEG networks is a subgraph which encompasses the strongest edges within the connectivity matrices, and reaches all nodes without creating cycling paths (16). Here the edges of the network are represented by their connectivity strength given by the PLI scores and the nodes of the network are the electrodes. The Prim’s algorithm (45) was used to estimate the MST with one minus the PLI score as input. Then, 10 MST measures and their variability (mean and SD; thus providing a total of 20 measures) were estimated for each frequency and participant. These measures are:

*Maximum betweenness centrality (BC_max_)*: is a measure of hubness or centrality (46) and it is proportional to the number of shortest paths that crosses a node in the tree (47). A node with high BC indicates high reachability of that node throughout the network. BC_max_ then represents the node with highest BC in the MST (48), and decreased BC_max_ suggests that the MST has decreased global efficiency (16).

*Diameter:* This is the largest distance or number of edges between any two nodes in the tree. An increased diameter indicates decreased network efficiency (23).

*Eccentricity:* Is the maximum distance or number of edges between a node and any other node in the tree. Here we summarised this measure as the average eccentricity across all nodes in the tree (48). Increased eccentricity in the MST suggests decreased network efficiency.

*Radius:* The smallest node eccentricity in the tree.

*Maximum degree (degree_max_)*: The degree of a node equals the number of edges connected to that node. The degree_max_ is the highest degree in the MST and belongs to the most connected node in the network (48). This is a measure of focused synchronisation, and the highest possible degree_max_ score would correspond to a MST that has a “star” topology (16) with only one central node connected.

*Leaf ratio:* The number of nodes connected to the MST by only one edge divided by the maximum number of possible leaves (M-1), where M equals 128 electrode nodes in this investigation.

*PLI mean:* The mean PLI score across the entire MST network.

*PLI leaf:* The mean PLI for all the edges connected to the leaf nodes.

*PLI root:* The mean PLI for the root node of the MST. Here the root is defined as node with the highest degree. Hence, PLI root is the mean PLI score for the edges connected to the node with highest degree in the MST.

*PLI height:* The difference between PLI root minus PLI leaf. PLI root equals the PLI leaf score if the MST has a star configuration (16).

MST measures were computed using functions from the Brain Connectivity (49) and the Network Analysis (50) toolboxes, as well as in-house functions in MATLAB.

### Statistics

Statistical tests for the clinical and demographic variables were implemented in SPSS (Statistical Package for the Social Sciences v22, IBM) and these are defined in Table 1. All EEG scores of variability, standard deviations and DFV, were log-transformed to approximate their distribution to a Gaussian. Differences in DF and DFV were investigated with a one-way ANOVA for the four groups and post-hoc unpaired t-tests for between group comparisons. Significance of MST measures was assessed in three steps. First, significant differences were investigated with two-way four-group ANOVAs for each MST measure as dependant variable and the following factors as independent variables: group (HC, AD, DLB, and PDD), frequency (delta, theta, high-theta, alpha, beta, and DF) and their interaction (group and frequency). Null hypotheses were rejected if either the diagnosis or the interaction effects were significant at a p-value<0.0025 thus accounting for the 20 MST measures assessed, i.e. Bonferroni correction. Secondly, for the significant two-way ANOVAs, post-hoc multiple one-way ANOVAs at each frequency band were evaluated and their significance was again Bonferroni corrected by the number of post-hoc tests, leading to an uncorrected p-value threshold of < 0.0007 (since 12 two-way ANOVAs survived the first stage of tests). Finally, group differences were assessed with multiple unpaired t-tests. For this latter step we were interested in three exploratory diagnostic scenarios of significant difference: AD vs DLB, DLB vs PDD and HC vs both AD and DLB (AD+DLB). Notice that differences in DF is a hypothesized test based on previous literature and as such the post-hoc tests of the one-way ANOVA are uncorrected, while the two-way four-group ANOVA tests for the multiple network measures are corrected for multiple tests or comparisons.

In order to assess relations of network measures with the clinical variables in our dementia patients, we implemented a multiple linear regression for the three dementia groups with clinical variable as the dependant variable and the network measure as the independent variable. Additionally, we included two dichotomous regressors for the DLB and PDD groups, in order to account for the group effects. The AD group was left as the reference group for being the largest group in our investigation. Before regression analysis, all measures of variability including DFV were log-transformed to approximate their distribution to Gaussian. The regression was considered significant if the first coefficient of the model (β_1_ for the network measure) was significant at a p-value < 0.05.

To assess the diagnostic utility of the studied measures in the three diagnostic scenarios, we implemented a logistic regression classification; sensitivity, specificity and area under the curve (AUC) were computed for each scenario, with confidence intervals estimated by statistical bootstrapping with 5000 iterations in Matlab. For the DLB vs AD case, scores from the PLI mean, PLI leaf mean, PLI root mean in the high-theta band, PLI mean, PLI root mean, PLI height mean in the beta band, DF and DFV were used as regressors. For DLB vs PDD, scores from the PLI mean, PLI root mean and PLI height mean in the alpha band were used in the logistic regression. The HC vs AD+DLB diagnostic scenario used twelve network features as regressors which also showed significant differences, Figure 5-Table Supplement 1. Diagnostic classification of HC vs PDD was not implemented. Additionally, in order to find the most significant discriminants for the three diagnostic scenarios a stepwise multiple linear regression was implemented.

### Network display and layout

In order to observe network topology characteristics within our studied groups, participants at the group median for the leaf ratio mean in the alpha band were selected as representative of each group. Leaf ratio mean scores were chosen for being consistently higher in the HC group compared with the dementia groups but not between the dementias. For each selected participant, the 49 PLI connectivity matrices spanning 50 seconds of EEG were averaged. From the mean connectivity matrix, the MST was extracted and displayed using a forced-directed graph layout (50), which facilitates graph visualisation by minimising edge crossings. This procedure was also repeated for the dominant frequency band for the same participants.

## Acknowledgments

The research was supported by the Northumberland Tyne & Wear Foundation Trust and the National Institute of Health Research Biomedical Research Centre (NIHR-BRC) at Newcastle University. The study was funded by an intermediate clinical Wellcome Trust Fellowship (WT088441MA) to J-P.T.

## Author contributions

LR.P executed the project, performed statistical analysis, and wrote the paper; R. C and S.G. executed the project, and reviewed the paper; X. K analysed data and reviewed the paper; MJ. F contributed on the design of the experiments and reviewed the paper; A. K executed the study and reviewed the paper; AJ. T and JT. O’B performed research; J-P. T. Designed the study, performed research and reviewed the paper.

---

Figure 1-Figure Supplement 1. The 10-5 EEG system layout. The occipital electrodes used in the estimation of DF and DFV are shown in blue colour with PO and O prefixes.

Figure 2-Table Supplement 1. Two-way ANOVA results for the effects of group (HC, AD, DLB and PDD), frequency (delta, theta, high-theta, alpha, beta and dominant frequency) and the interaction. P-values shown adjusted for multiple tests. PLI height (SD) was not significant after Bonferroni correction.

Figure 2-Table Supplement 2. Global effects of group diagnosis; two-way ANOVA tests.

Figure 3-Table Supplement 1. Acronyms and meaning of network measures and other variables.

Figure 3-Table Supplement 2. Minimum spanning tree (MST) measures. Man and standard deviation (SD), Significant one-way four-group ANOVAs with p-value < 0.05, Bonferroni corrected for multiple tests.

Figure 5-Table Supplement 1. Network measures and dominant frequency variables used for the logistic.

Table 2-Supplemental Table 1. Multiple linear regressions for relations of network measures with clinical variables. Regression model: Clinical variable = β_1_*Network variable + β_2_*D_DLB_+ β_3_*D_PDD_ + Intercept.

